# Allosteric Control of Super-Agonism in a Ligand-Gated Ion Channel

**DOI:** 10.1101/2025.05.23.655750

**Authors:** Franco Viscarra, Teresa Minguez, Isabel Bermudez-Diaz, Philip C Biggin

**Author notes:** Senior author (Philip C Biggin). Bluesky: @philbiggin.bsky.social (Philip C Biggin).

## Abstract

Ligand-gated ion channels open in response to the binding of an agonist. The agonist binding site is typically located tens of Angstroms away from the channel gate, which lies within the membrane and thus the gating mechanism is considered a classic example of allostery within proteins. The multi-subunit nature of these proteins also means that modulatory effects on the gating process can also be mediated by several other distinct regions – so called allosteric modulatory sites. One of the most well-studied channels in this regard, is the nicotinic acetylcholine receptor. Super-agonists are compounds that can produce a greater maximal response than the endogenous ligand (acetylcholine in this case). They are able to stabilize the open state of the ion channel and this can have important consequences for neuronal signaling. Some super-agonist effects can be mediated through the orthosteric binding site, but others can be mediated by alternative binding sites. The latter are often much harder to identify and can depend very precisely on the subunit composition of the receptor. In this work we sought to identify the mechanism by which TC-2559, a known super-agonist that acts as such only at one particular combination of neuronal nicotinic acetylcholine receptors – the high sensitivity receptor which is comprised of 2 alpha subunits and 3 beta subunits (as opposed to the low sensitivity receptor which has 3 alpha and 2 beta subunits). By using advanced computational methods, supported by two-voltage electrode clamp experiments, we were able to show that TC-2559 not only binds to the orthosteric but also binds to the unique b2-b2 interface of the HS receptor. The binding of TC-2559 to this interface exerts unique interactions that other agonists are not able to make, but more importantly it induces changes in the interface that support that concept of an allosteric gain in overall efficacy. Our results highlight how allosteric control exists to modulate receptor function.

## INTRODUCTION

Allostery, the process by which a binding event at one site of a macromolecule influences activity at a distant functional site, is a cornerstone of biological regulation, enabling control over processes like signal transduction, metabolism, and gene expression.^1^ The nicotinic acetylcholine receptor (nAChR) is the quintessential example of an allosteric protein^2^ and indeed the analysis and supporting theory has been substantially developed over the years. This is supported by web-resources that collate structure-function relationships in the context of allosteric signalling within the nicotinic acetylcholine receptors.^3^ NAChRs are pentameric ligand-gated ion channels that converts neurotransmitter binding into electrical signals across synapses. In recent years, cryo-EM structures and molecular simulations have provided deeper insight into how binding of ligands^4-6^ or indeed lipid composition,^7-11^ shifts the conformation of the receptor between three key states: resting, active (open) and desensentized. The presence of multiple binding sites and conformational equilibria had made these receptors very much a paradigm for understanding allosteric communication within proteins.

NAChRs most commonly assemble as heteromers and can do so with variable stochiometries^12^ The α4β2 nicotinic acetylcholine receptor (nAChR) is the predominant nicotinic acetylcholine receptor in the mammalian brain, making up 90% of high-affinity nAChRs and serving as the primary nicotine-binding subtype.^13^ This makes it the main target for the treatment of nicotine addiction^14,15^ (with partial agonists such as varenicline^16^) and a potential target for a wide array of central nervous system disorders.^15,17-19^ This receptor assembles in two stoichiometries: (α4)3(β2)2, known as low-sensitivity (LS), and (α4)2(β2)3, known as high-sensitivity (HS)^20^ (**Fig. 1**). The orthosteric agonist binding site at the two stoichiometries of the α4β2 nAChR are structurally identical, leading to similar agonist sensitivity, albeit with a range of efficacies.^15^ The differences in agonist efficacy can be accounted for partly by the additional agonist site at the α4-α4 subunit interface of the LS receptor. This additional site generally increases the efficacy of agonists,^15^ with the notable exception of agonists TC-2559 (**Fig. 1**) and Sazetidine-A. Neither of these ligands can access the agonist site at the α4-α4 interface of the LS receptor, where they display partial agonism.^21^ In contrast, at the HS receptor, TC-2559 behaves as a super-agonist (with its maximum response being almost 4-fold greater than acetylcholine)^15,22^ and Sazetidine-A as a full agonist.^23^

**Figure 1.**
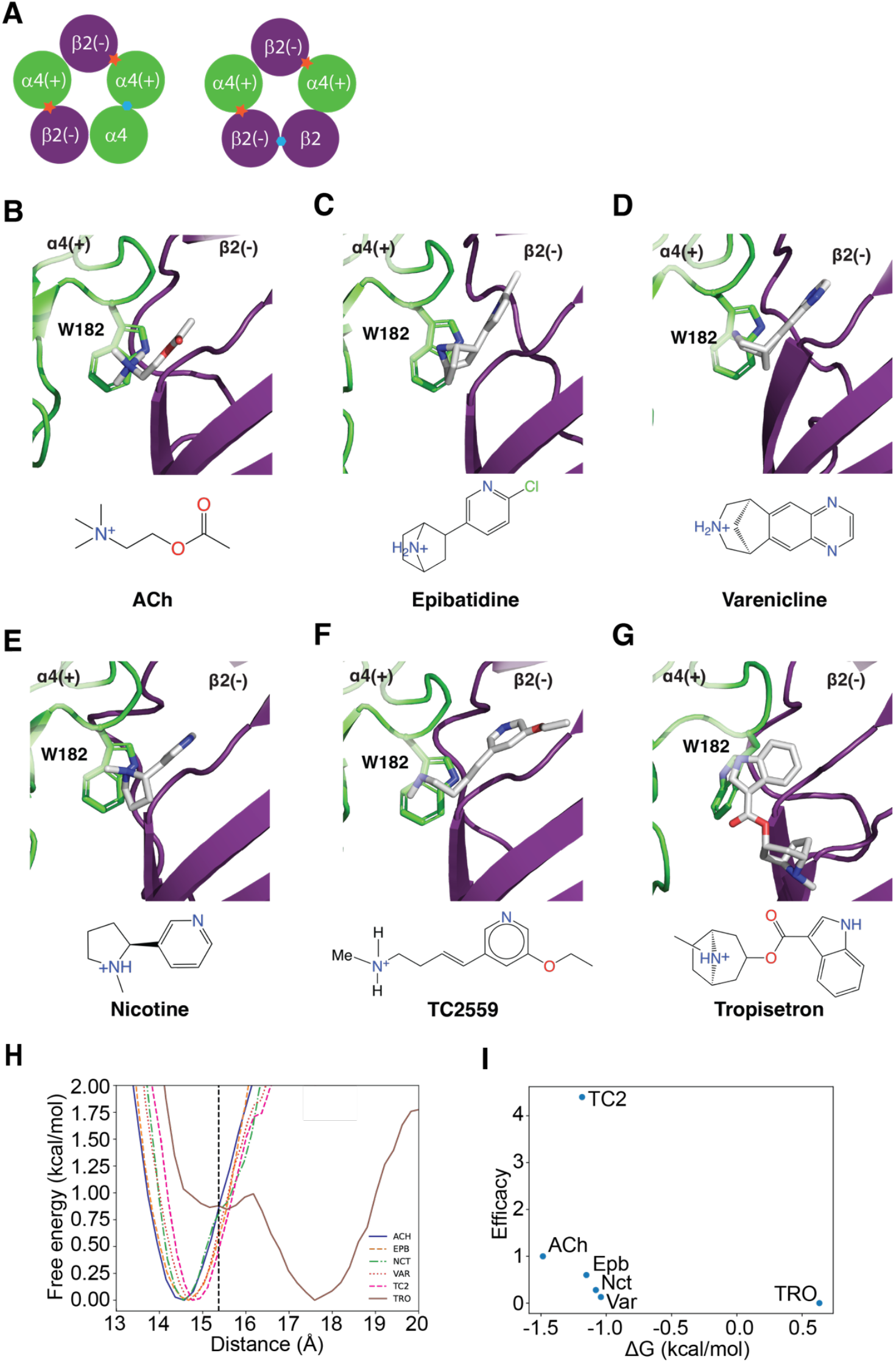
α4β2 stochiometries and agonist or antagonist binding at the orthosteric site. (A) Cartoon representation of the low (α4)_3_(β2)_2_ and high (α4)_2_(β2)_3_ sensitivity stoichiometries of the α4β2 subtype nAChR. The location of the orthosteric and unique unorthodox binding sites are highlighted with orange stars and blue hexagons respectively. (B-G) Clustered poses from REST simulations of selected agonists (B-F) and the antagonist tropisetron (G) at the α4β2 interface (orthodox binding site). The α4 subunit is coloured green, the β2 subunit is coloured purple, and the ligand is coloured in silver. W182, key residue of the binding site is shown in stick representation. Ligands are drawn assuming pH = 7 with pKa predictions from https://xundrug.cn/molgpka/silicopka. Loop C is removed for clarity. (H) Free energy landscape of loop C capping from REST simulations. A. free energy landscape over the loop C distance to the protein COM. The dashed line represents the value of the Cryo-EM structure (6CNJ). (I) Energy difference between the activated and inactive states plotted against the agonist efficacy (assumed 0 for the antagonist, Tropisetron).

Since the orthosteric agonist binding sites located at the α4(+)-β2(-) interfaces are structurally identical, they are unlikely to confer receptor stoichiometry-specific functional signatures. However, because the fifth subunit in the α4β2 can be α4 or β2, each α4β2 nAChR stoichiometry has a unique subunit interface, an α4(+)-α4(-) binding interface at the LS receptor and a β2(+)-β2(-) interface at the HS. This suggests that the efficacy of TC-2559 or Sazetidine-A at the HS receptor may be driven by binding to β2-β2 interface.

In the LS stoichiometry, the α4-α4 site harbours a modulatory site that seems to be homologous to the benzodiazepine site in the GABAA receptor.^24-27^ Thus, by analogy to the LS and GABAA receptors, it is reasonable to suggest that TC-2559 or Sazetidine-A may display higher efficacies at the HS by binding to both the orthosteric agonist sites and a region in the β2-β2 interface. Binding to the β2-β2 interface may potentiate the functional effects of the agonists mediated by binding to the orthosteric sites.

In this work, using TC-2559 as a model ligand, computational approaches, mutagenesis, and functional assays we explored the view that the β2-β2 site may harbour an allosteric binding site that modifies the efficacy of agonists bound at the orthosteric agonist site of the HS receptor.

## Results

### Do agonists of the HS receptor behave similarly in the orthosteric site

There is currently structural information only for nicotine, acetylcholine, and varenicline bound to α4β2 nAChR.^28,29^ Thus, in the first instance we investigated the binding of TC-2559 to the orthosteric site and compared it with other well-characterized agonists that exhibit a range of efficacies (acetylcholine, epibatidine, varenicline, nicotine and the antagonist, tropisetron (**Fig. 1B-G**). To do this, we used replica-exchange solute tempering (REST) simulations (see Methods for details) to explore binding modes. Clustering of the trajectories rendered similar binding modes for the tested agonists, with the charged amino group of the agonists interacting with W182 (**Fig. 1B-F**).

Loop C, which forms a cap across the front of the binding site and is a well-conserved feature of all pentameric cys-loop receptors. The extent of how tightly this capping process occurs has previously been suggested to correlate with agonist efficacy and thus we were interested to know if TC-2559, a super-agonist, would reveal any difference in this aspect. We determined the extent of capping but using the distance between the centre of mass (COM) of residues in the tip of loop C (residues 223-226) and the COM of the interface. The free energy surface (FES) over this collective variable (CV) was calculated with the Boltzmann relationship for each ligand. The FES separated the agonists (acetylcholine (ACh), epibatidine, nicotine, varenicline, and TC-2559) from the antagonist (tropisetron), with the basins of the agonists lying to the left in a more capped conformation and the basin of tropisetron in a more open one (**Fig. 1H)**, as expected based on previous works ^5,6,30,31^. In this FES, the active state was defined as having a CV value lower than 15.37 Å (the CV value of the starting structure: 6CNJ), and the inactive as having a value higher than 15.37 Å. The free energy difference between the Boltzmann averages of the active and inactive states shows a relationship with the efficacy of the agonist (**Fig. 1I**). Interestingly, TC-2559, a ligand that behaves as a superagonist at HS α4β2 nAChR, does not comply with this pattern, having a lower free energy difference between states than ACh, a full agonist. This finding supports the view that TC-2559 exerts its super-agonism at the HS receptor through a mechanism that does not involve binding to the orthosteric agonist site.

### Aspartates at β2 Loop C and Loop F are Critical for TC-2559 Function

Since the β2-β2 interface is unique to the HS receptor, (and TC-2559 only exerts its super-agonist effect at the HS), binding of TC-2559 to this β2-β2 interface was also explored by performing REST simulations. These simulations revealed that the charged amine of TC-2559 had strong interactions with the carboxyl oxygens of D195 and D196 of Loop F and D217 and D218 of Loop C. These interactions are not present for ACh, nicotine, epibatidine or varenicline (**Fig. 2A**). Alanine substitution of these residues and subsequent two-electrode voltage clamp experiments demonstrated that altering these residues greatly diminished the efficacy and potency of TC-2559 without affecting the ACh response (**Fig. 3A and 3B**).

**Figure 2.**
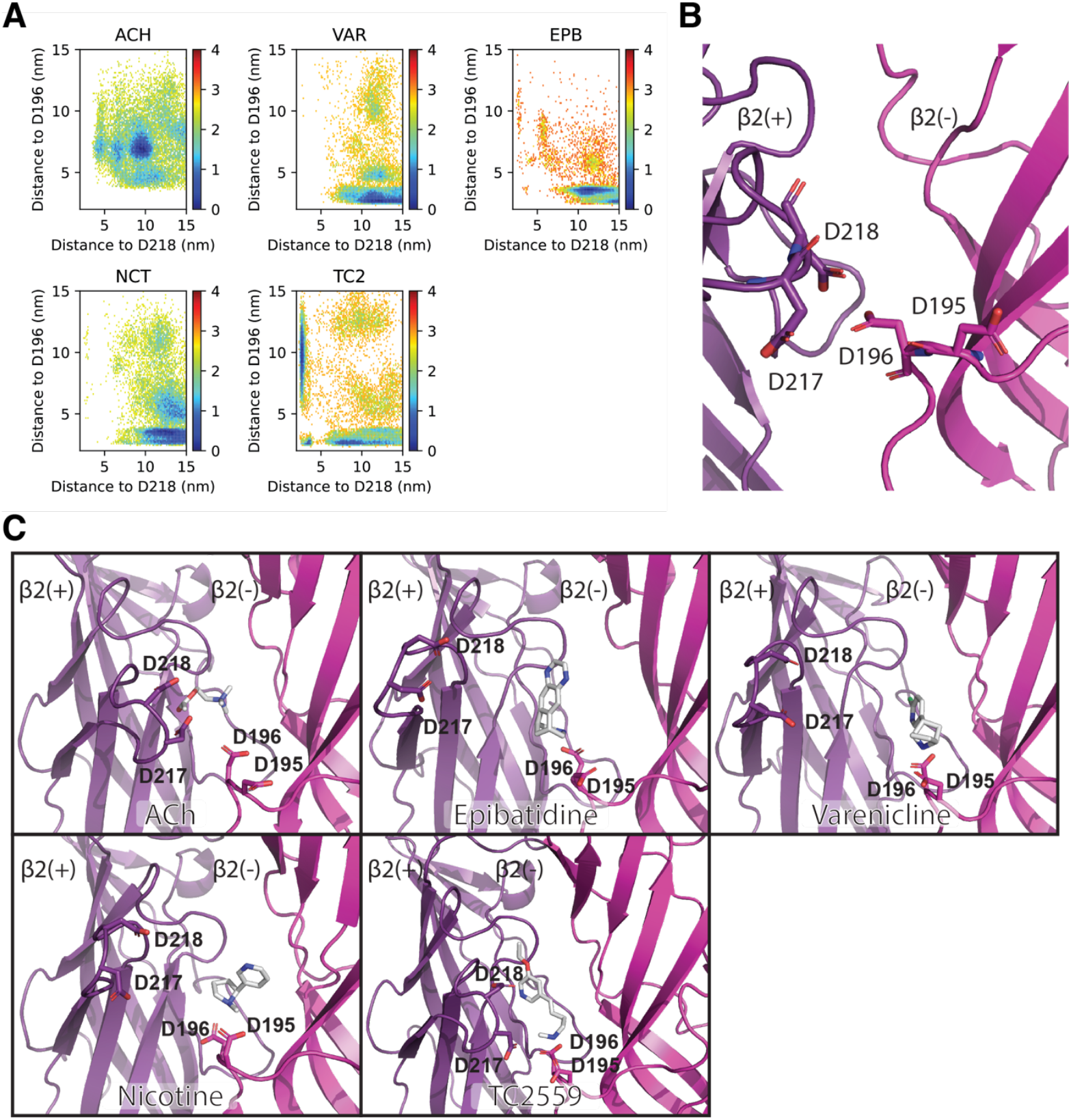
REST simulations of agonists at the β2-β2 interface. (A) 2-dimensional FES for the interaction of the charged amino group of every agonist with the charged oxygen of D196 and D218. (B) Aspartates in the β2(+) (purple)/β2(-) (magenta) interface. (C) Clustered conformations obtained from REST simulations. The β2(+) interface is coloured purple, the β2(-) is magenta, and the ligand is coloured grey. The key aspartates and the ligand are shown in stick representations. TC-2559 is the only compound that interacts with both D196 and D218.

**Figure 3.**
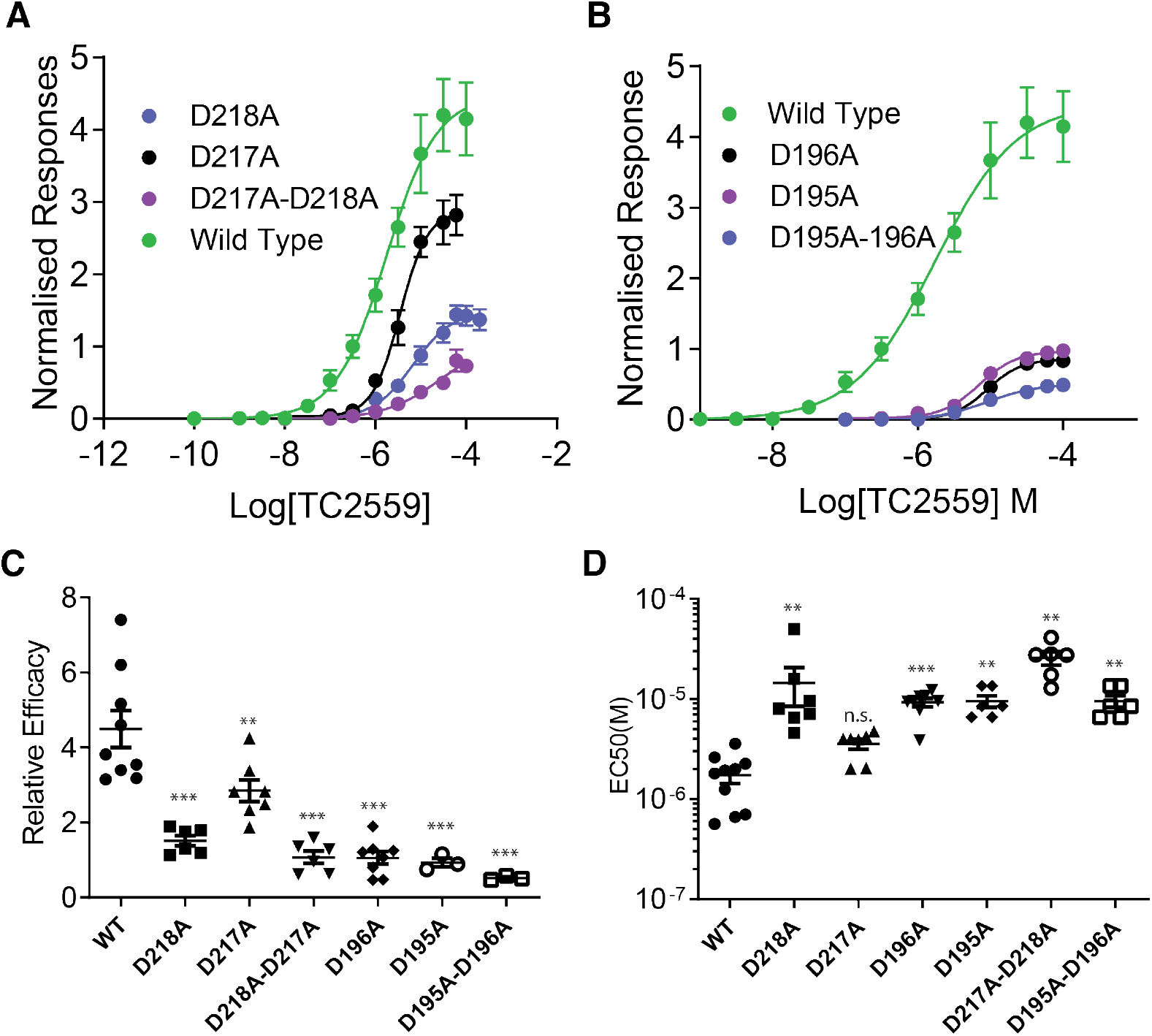
Two-electrode voltage clamp experiments of β2-β2 interface mutants. (A) Concentration-response curves of TC-2559 in the loop C mutants. (B) Concentration-response curves of TC-2559 in the loop F mutants. (C) Efficacy of TC-2559 on each mutant compared against the wild-type. (D) Affinity of TC-2559 on each mutant compared against the wild-type. Statistical significance was tested using a one-way ANOVA with multiple comparison tests: ** = p < 0.01; *** = p < 0.001; n.s. = not significant.

D217A significantly reduced the efficacy of TC-2559 (2.8 ± 0.8) but kept its super-agonist profile. D218A produced a more dramatic decrease in efficacy (1.5 ± 0.3), similar to the effect observed in the presence of both mutations (D217A/D218A) (1.1 ± 0.4) (**Fig. 3C**). In the case of the loop F aspartates, both D195A or D196A eliminated the super-agonism (D195A efficacy: 0.93 ± 0.2; D196A efficacy: 1.1 ± 0.5) with the presence of both mutations rendering TC-2559 into a partial agonist (0.52 ± 0.1) (**Fig. 3C**). Interestingly, all the mutants significantly decreased the affinity of TC-2559, except for D217A (**Fig. 3D**).

### Displacement of β2 Loop B is Induced by TC-2559

Having established the binding mode and key interaction partners in the allosteric binding site, we next sought to try and understand how TC-2559 confers its super-agonist effect. To that end, we used funnel metadynamics (FunMetaD) simulations to try and gain information about the actual binding event. The ligand was simulated in the β2-β2 wild-type (WT), D217A-D218A and D195A-D196A receptors for a total of 25 µs each. From these simulations, and to our surprise, no significant difference in the binding free energy of the WT compared with the mutants was observed. However, the FES of ligand binding showed a shift in the binding mode of TC-2559 with the low energy basins located in different sections of the interface when the aspartates are mutated (**Fig. 4A)**. Representative structures extracted from clustering the basins show that upon binding TC-2559 establishes simultaneous interactions with loops B and F, causing significant Loop B displacement (**Fig. 4B**).

**Figure 4.**
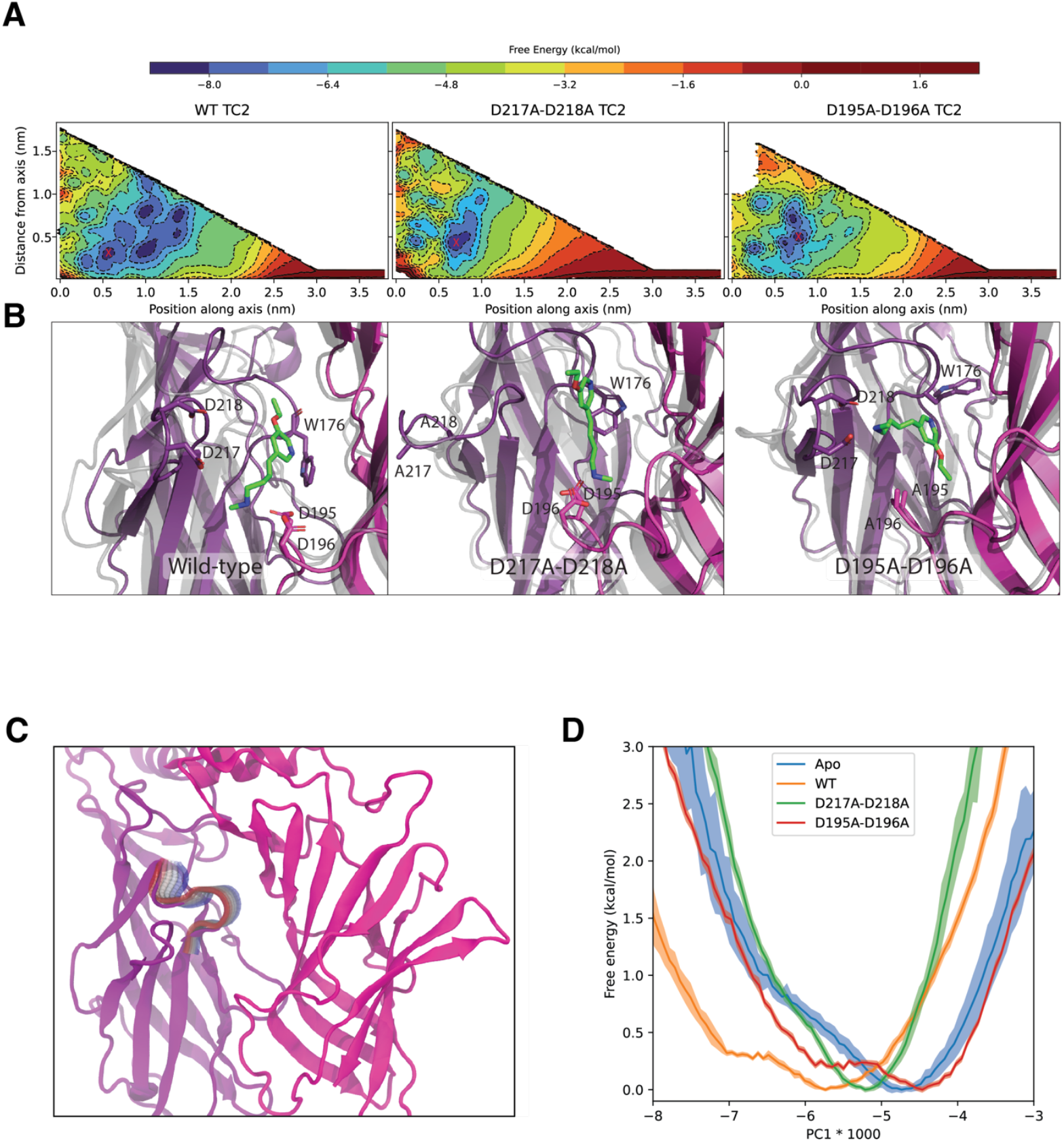
Funnel metadynamics of TC-2559 at the β1-β2 interface. (A) Free energy landscape of the ligand COM along the funnel coordinates (position along the funnel axis and distance from the funnel axis) for the wild-type and mutants. The minimum energy basins are highlighted with a red X. (B) Representative structures of each basin are obtained through RMSD-based clustering of the trajectory frames in each basin. The β2(+) subunit is coloured in purple, while the β2(-) subunit is coloured in magenta. The Cryo-EM structure (6CNJ) is superimposed in light grey. Residues W176, D195, D196, D217, and D218 are shown in stick representation. TC-2559 is coloured green. (C) Schematic representation of the motion described by PC 1 at the β2-β2 interface. The β2(+) subunit is coloured in purple, while the β2(-) subunit is coloured in magenta. The alpha carbons included in the calculation are coloured on a scale from red to blue going from the apo-like to the bound-like conformations. (D) FES of PC1 for the apo and TC-2559 with WT, D217A-D218A, and D195A-D196A receptors. The shaded areas represent the standard deviation calculated with 1000 bootstraps.

To characterise this conformational change, the PC of the Loop B Cα (residues 174-182) between the frames of the bound basin from the WT simulation and an apo simulation of the β2-β2 WT interface (10 replicas of 500 ns each) were calculated (**Fig. 4C**). The FES was projected over the first PC (**Fig. 4D**) for the WT and mutant receptors. The WT simulation in the presence of TC-2559 has a minimum energy at a PC value of -5.7, while the apo simulation is most stable at -4.7. In the mutant simulations, D217A-D218A elicited a smaller deviation from the apo state, with a minimum energy state in the -5.2 and D195A-D196A did not induce a deviation towards a WT-like state with a global minimum at -4.5.

## Discussion

The agonist conformations obtained from MD simulations performed at the α4-β2 interface concur with previous observations reported in the literature, validating the REST technique in these conditions. First, the binding mode of both nicotine and ACh aligns closely with the binding poses of these ligands in Cryo-EM structures of the α4β2 nAChR^28,29^ (**Fig. S1**). Moreover, the displacement of Loop C has been well established as a conformational change occurring upon agonist binding. Early studies have shown that agonists, but not competitive antagonists, protect the binding site of nAChRs against reduction from dithiothreitol in *Torpedo californica* electric membrane, suggesting a “closing” conformational change in the binding site upon agonist binding.^32^ Later, crystal structures of AChBP from *Aplysia californica* display Loop C capping in the presence of epibatidine, while in the apo and methyllycaconitine-bound structures, Loop C appears in a more open fashion.^30^ More recent structures resolved from Cys-loop receptors, such as the GluCl^31^ and the α7 nAChR,^5,6^ also show this conformational change in the presence of activating agonists. Additionally, loop C truncation eliminates ACh activation of nAChR, although it does not affect unliganded gating.^33^

Allosteric gating models, such as the one described by Plested, postulate that a transition between low-affinity and high-affinity binding domain conformations (the latter characterised by the capping of Loop C) is a favourable step in agonist-induced gating.^34^ Furthermore, the efficacy of a ligand is related to the binding energy of the ligand to these two states.^35^ Thus, theoretically, it is possible to estimate the efficacy of a ligand by calculating the difference between the absolute binding free energy of the ligand to the protein in each conformation^36^ or, alternatively, by calculating the free energy difference between the states in the presence of the ligand. This work uses these ideas to relate Loop C’s displacement from REST simulations in the presence of different ligands to their relative efficacy. However, this comes with limitations: since it is necessary to apply a flat-bottom restraint potential on the ligand to avoid dissociation at high-temperature replicas,^37^ this may affect the ligand’s ability to induce conformational changes, adding unknown biases to the free energy landscape.

Nevertheless, the free energy difference between the capped and uncapped conformations in the REST simulations show a direct relationship with ligand efficacy, with the antagonist stabilizing a non-capped conformation. It might be expected for a compound like TC-2559, whose efficacy surpasses that of ACh, that it would stabilize an activated-like conformation even further than the endogenous neurotransmitter. Calculations show this is not the case, and simulations with TC-2559 show a behaviour similar to the partial agonists studied here (**Fig. 2**). Therefore, our evidence suggests that TC-2559 does not exert its signature super-agonism through binding at the orthosteric binding site. Taken together with the stoichiometric selectivity of this effect, we propose that, in addition to its agonist binding site at the α-β interface, TC-2559 binds to a positive allosteric site at the β2-β2 subunit interface, which is unique to the HS receptor.

In the LS stoichiometry of the α4β2 receptor, binding of ligands to the unorthodox α4-α4 binding site enhances the probability of opening of the receptor, allowing partial agonists, such as cytisine and varenicline,^15,21^ to display higher efficacy than at the HS receptor. Furthermore, epibatidine, a partial agonist at the HS stoichiometry, evokes a response even higher than ACh’s in the LS receptor.^15^ Additionally, the α4-α4 interface can bind positive modulators like NS9283 and CMPI11,25, although these modulators enhance the affinity of orthosteric ligands rather than their efficacy. At the HS stoichiometry, binding to the β2-β2 site could generate a similar effect. However, a direct comparison is difficult because the positive site at the α4-α4 interface is formed by the α4 subunit, which includes key structural features essential for agonist-induced gating—specifically, a longer Loop C and an electron-rich aromatic box.

In contrast, while the β2-β2 subunit interface largely preserves the aromatic ring, it cannot form the crucial π-cation interaction with the agonist due to the presence of R175. In the α4 subunit, this position is occupied by a glycine instead. R175, positioned at the base of the binding site, carries a positive charge that mimics a positively charged agonist. It is sandwiched between tyrosine residues from Loop C and Loop A, effectively compensating for the electron-rich π pocket. This rearrangement also causes a shift in loop B’s tryptophan, which, in Cryo-EM structures, appears to be oriented away from the binding site (REF – is this Walsh?)

REST simulations of TC-2559 in the β2-β2 interface confirm these observations by suggesting that TC-2559’s positively charged amino group interacts with negatively charged aspartates in Loop C and Loop F of the β2 subunit rather than with the aromatic box as in the orthosteric binding site. The importance of these aspartates is confirmed by alanine substitution of these amino acids, which significantly affects both the affinity and efficacy of TC-2559. It is worth noting that although Loop C’s aspartates are unique to the non-canonical β2-β2 and β2-α4 interfaces, the loop F’s aspartates are also present in the orthosteric α4-β2 site. Still, the current evidence suggests that at the orthosteric binding site of Cys-loop receptors, this loop plays a role in ligand accommodation and selectivity rather than receptor activation,^6,38^ therefore its alteration should not produce a noticeable effect on ligand efficacy, yet the mutants of loop F’s aspartates produces a significant decrease in TC-2559’s efficacy (**Fig. 3C**). Additionally, the substitution of loop F’s aspartates does not affect the pharmacology of ACh (**Fig. S2**).

Metadynamics simulations of TC-2559 in the β2-β2 binding site also highlighted the importance of these aspartates, suggesting that they are key for determining TC-2559’s binding mode and the conformational changes induced by this ligand. Notably, a displacement of Loop B is induced upon binding of TC-2559 to this site. This loop contains the key conserved tryptophan analogous to the one that engages in cation-π interactions with agonists at the α4 subunit of the orthosteric binding site. Furthermore, Cryo-EM structures from the 5-HT_3A_ receptor and the α7 nAChR have shown a displaced Loop B in the agonist-bound structures compared to the apo.^6,39^ Loop B is directly connected to the coupling’s region Cys-loop through the β7 sheet, which links the motions of Loop B with the transmembrane domain. Moreover, unique to the β2 subunit, there is a salt-bridge connecting K160 of the Cys-loop (a threonine in the α4 sequence) to D293 in the M2-M3 linker (a valine in the α4). This provides for a putative allosteric communication pathway between the motions induced by TC-2559’s binding and a displacement of the pore lining the M2 helix, which could result in further stabilization of the open state and the observed super-agonism at a macroscopic level. Additionally, computational studies have highlighted the role of β2’s Loop F in allosteric communication between the extracellular and transmembrane domains upon nicotine binding.^40^

Another parallel to a similar binding site is the benzodiazepine site in GABA_A receptors. This site is located at the interface between the positive face of an α subunit and the negative face of a γ subunit. In GABA_A receptors, the α subunit typically contributes to the negative face of the orthosteric binding site, whereas the γ subunit does not participate in orthosteric ligand binding. As a result, this site can be considered partially analogous to the β2-β2 interface, as it also contains a non-orthosteric positive face. Mutational studies in GABA_A receptors have identified key residues involved in benzodiazepine binding. In the α subunit, amino acids in Loops C and B play a crucial role in benzodiazepine function, influencing both binding affinity and efficacy. In contrast, in the γ subunit, mutations in Loop F significantly reduce benzodiazepine potentiation. Notably, the D192A mutation completely abolishes diazepam potentiation, and this residue is positioned similarly to D196 in the β subunit’s loop F.

## Limitations of the study

To date, no experimentally resolved structure of the α4β2 nAChR bound to TC-2559 or any ligand bound to the β2-β2 interface has been reported. Thus, the work presented here is heavily based on *in-silico* structural models derived from simulations started from the available cryo-EM structures of the α4β2 nAChR bound to nicotine, which may not reflect the correct conformational state of a receptor bound to TC-2559. Enhanced sampling simulation methods were used to overcome this limitation; nevertheless, there are also limitations concerning their application. In the case of REST, to keep the number of replicas manageable, it is necessary to heat only a small portion of the system, which means that only the motions of the ligand and surrounding residues were accelerated, while the rest of the protein was kept with an unmodified Hamiltonian, enhancing the sampling of only local motions within the binding site. Also, to avoid ligand dissociation in high-temperature replicas, a flat-bottom restraint was applied to the ligand, which can influence the ability of the ligand to induce conformational changes in an unknown (and hence, unaccounted for) manner. Additionally, all simulations were performed in a truncated construct that contains only the extracellular portion of the relevant interface studied. Although the capacity of this construct to reconstruct native ligand poses has been validated, there are still potential artefacts introduced due to having normally buried sites solvent-exposed.

Experimentally, the mutations introduced to the complementary β2 face (Loop F D195A and D196A) are present both in the canonical α4-β2 site and the proposed β2-β2 site, making the interpretation of the results ambiguous because it is impossible to separate the effects over the two interfaces. Nevertheless, the evidence supports the idea that Loop F in the α4-β2 site influences ligand accommodation, and thus binding, rather than signal transduction, which should be reflected mostly on the ligand’s affinity rather than the efficacy, as observed in our data.

## OUTLOOK AND CONCLUSIONS

In summary, this work presents a combination of computational and experimental evidence that supports the hypothesis of TC-2559 exerting its super-agonist effect at the HS α4β2 nAChR through binding at a unique allosteric binding site. On the one hand, computational calculations at the orthosteric binding site show that TC-2559’s binding mode aligns with that of classical partial agonists. On the other hand, investigation of TC-2559’s binding at the β2-β2 site revealed key residues whose modification alters the ligand’s behaviour in both experimental and computational tests. Moreover, analogy with other modulators of Cys-loop receptors supports the possibility of positive modulation at a non-canonical extracellular binding site, mediated by the key structural motifs identified in this study. This binding site provides a target for the development of novel positive allosteric modulators selective for the HS stoichiometry of the α4β2 nAChR, which have not yet been reported. Due to their widespread distribution in the brain, the α4β2 nAChRs play a significant role in several diseases. They are implicated in nicotine addiction by mediating dopamine release and receptor desensitization^41^, in Alzheimer’s disease through their interaction with amyloid-beta peptides^42^ and in Parkinson’s disease by modulating dopaminergic pathways^43^. Additionally, α4β2 nAChRs are of interest in ADHD^44^, pain^45^, and depression^46^. Involvement of nAChRs in neurodegenerative diseases and mood disorders is mainly supported by epidemiological observations. These studies correlate disease onset or symptom relief with smoking ^43,47^, highlighting the relevance of HS α4β2 nAChRs, since they are likely the main receptor type activated by nicotine at physiological concentrations in the brain when smoking^48,49^. Additionally, studies have demonstrated the analgesic effect of Sazetidine-A, a selective agonist of the HS stoichiometry of the α4β2 nAChR^50^, further supporting the importance of HS α4β2 nAChRs in pain regulation. Finally, although a putative allosteric mechanism has been proposed, more studies are required to confirm the exact mechanisms and pathways.

## RESOURCE AVAILABILITY

### Lead contact

Further information and requests for resources and reagents should be directed to and will be fulfilled by the lead contact.

### Data and code availability

Simulations and structure files have been deposited at Zenodo at https://doi.org/10.5281/zenodo.15387367 and are publicly available as of the date of publication.

## ACKNOWLEDGMENTS

F.V. was supported by Becas Chile for overseas PhD studentship. IB-D and FV also thanks the Oxford Brookes University Research Excellence funding scheme. Computing was supported via the EPSRC ARCHER2 UK National Supercomputing Service granted via the High-End Computing Consortium for Biomolecular Simulation, (HECBioSim -http://www.hecbiosim.ac.uk), supported by EPSRC (EP/L000253/1).

## AUTHOR CONTRIBUTIONS

Conceptualization, FV, IB-D and PCB; methodology, all authors; Investigation, FV and TM; writing—original draft, FV; writing—review & editing, all authors; funding acquisition, FV, IB-D and PCB; resources, IB and PCB; supervision, IB and PCB.

## SUPPLEMENTAL INFORMATION

Document S1. Figures S1–S2.

## METHOD DETAIL

### Molecular Dynamics Simulations

#### System Preparation

The coordinates for the HS α4β2 nAChR were obtained from the Cryo-EM structure PDB ID: 6CNJ.^29^ A construct consisting of the truncated extracellular portion of the α 4-β2 or β2-β2 interface, as appropriate, was prepared. Harmonic restraints were applied to the five initial and final residues of each chain to compensate for the absence of the other subunits. This type of construct for simulating binding sites of the α4β2 nAChR has been successfully validated in previous works.^24^ The protein was parameterized using Amber99SB force field.^51^ The ligand structure was produced with RDKit,^52^ and the parameters were generated with AmberTools22^40^ using the GAFF force field.^53^ The ligand was inserted in the binding site pocket by selecting the first conformation generated by AutoDock Vina.^54^ Then, the simulation box was solvated with TIP3P water^55^ and neutralized with 0.15 of NaCl. The system was energy minimized and then equilibrated for 1 ns in NVT and then NPT ensembles. All simulations were performed with GROMACS 2022^56^ patched with PLUMED version 2.8.0.^57^

#### REST Simulations

REST simulations were run in the α4-β2 and the β2-β2 interface. For the α4-β2 interface the solute region was defined as Y126, W182, T183, Y223, C225, C226 and Y230 of the main (+) chain and W82, V136, F144 and L146 of the complementary (-) chain. In the case of the β2-β2 interface the solute region was defined as Y120, S175, W176, T177, D217, D218 and Y221 of the main (+) chain and W82, V136, F144 and L146 of the complementary (-) chain. In both cases, the ligand was kept in the binding site using a flat-bottom restraint at 10 and 8 Å from the binding site COM, respectively. The system was simulated in 16 replicas with temperatures ranging from 310 to 1000 K with exchanges between replicas being attempted every 1000 steps. Representative conformations from the target temperature replica (310 K) were extracted by clustering using the GROMOS algorithm.^58^

#### Funnel Metadynamics

FunMetaD simulations were run at the β2-β2 interface with the WT, β2 (+) D217A-D218A or β2 (-) D195A-D196A mutants. The funnel was set with the axis between the centre of the binding site and the apex of loop C, with a z_cc_ of 30 Å, an α of 0.5, and a cylinder radius R_cyl_ of 1 Å.^59,60^ The CVs biased by the metadynamics were the position of the ligand COM along the funnel axis (fps.lp) and the distance of the ligand from this axis (fps.ld). These CVs were biased with a Gaussian width of 0.02, and an initial height of 2 kJ/mol. The free energy surface of the FunMetaD simulations over any arbitrary CV was calculated by calculating the weight of each frame using the expression 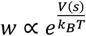, with *V*(*s*) being the metadynamics bias potential, and then performing a weighted histogram.^61^

#### Two-Electrode Voltage-Clamp Recordings

Concentration response curves (CRCs) of TC-2559 and ACh were constructed by normalizing the current responses to the maximum response elicited by ACh and fitting them to a Hill equation. The responses were recorded using two-electrode voltage-clamp recording at a holding potential of -60 mV using an Oocyte Clamp OC-725C amplifier (Warner Instruments, USA) or the automated HiClamp system (Multi Channel Systems, Reutlingen, Germany), as previously reported.^62^

#### Site-Directed Mutagenesis

The β2 mutants were produced using the Q5 Site-Directed Mutagenesis kit from New England Biolabs (U.K.). The presence of the mutation and the absence of unwanted mutations were confirmed by sequencing the entire cDNA insert (Eurofins, U.K.).

## QUANTIFICATION AND STATISTICAL ANALYSIS

Nonlinear fitting of the CRCs and the statistical analyses of efficacy and potency were carried out with GraphPad Prism 8 (GraphPad, USA). Unless otherwise stated, values are reported as the mean and dispersion as the SEM. Analysis of MD trajectories was performed using in-house Python scripts based on the MDAnalysis package.^63^ PCA of the trajectories was performed using the gmx covar tool and calculated for all trajectory frames with GROMACS^43^ patched with PLUMED. ^44^ Molecular graphic figures were generated using PyMOL,^64^ except for the principal component graphic, which was generated using VMD.^65^

In electrophysiological experiments (Figs. 3 A-D) every CRC was constructed from 6-10 oocytes from at least two different *Xenopus* frogs. Peak currents were normalised to the maximal ACh response obtained in the same cell, pooled, and fitted to the four-parameter Hill equation. Shapiro-Wilk tests showed normal distributions for all efficacy but not for all potency groups. Thus, efficacy estimates were compared among constructs with a one-way ANOVA followed by Dunnett’s post-hoc test against the WT receptor, while potency estimates were compared with a Kruskal-Wallis test followed by Dunn’s multiple-comparison test versus WT. Statistical significance was defined as P < 0.05, and exact P values are reported in the figure panels (*** P < 0.001; ** P < 0.01; n.s., not significant).

