## Supplemental Information for "Allosteric Control of Super-Agonism in a Ligand-Gated Ion Channel"

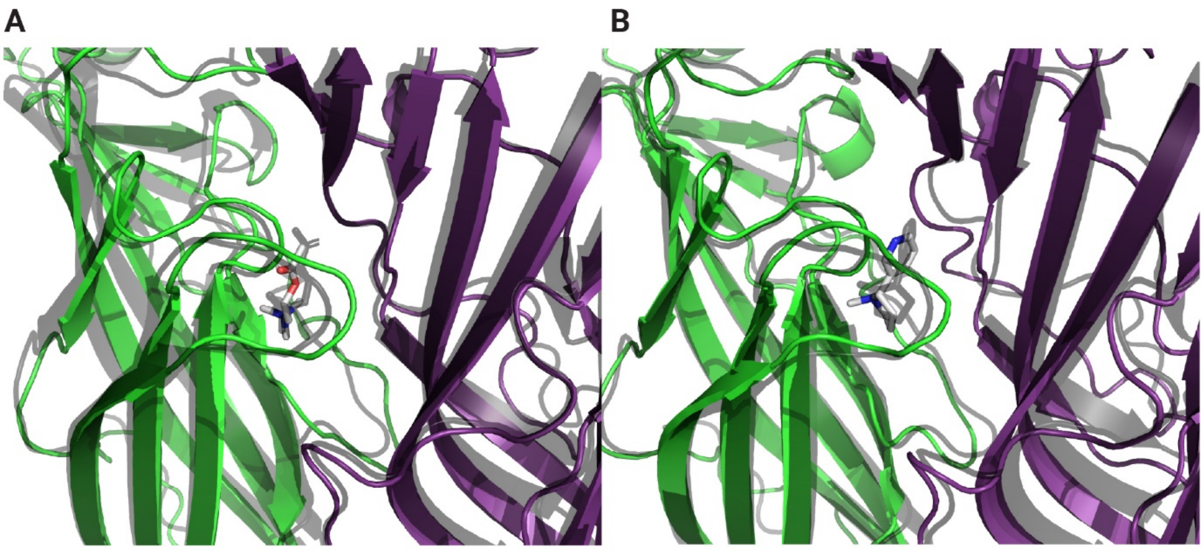


**Figure S1. Clusters from REST simulations superimposed to Cryo-EM structures.**

*(**A**) Best cluster superimposed with one of the α4-β2 interfaces with nicotine bound between the α4 (green) and β2 (purple) interface from RCSB:6CNJ.*

*(**B**) Best cluster superimposed with one of the α4-β2 interfaces with ACh bound from RCSB:8ST4. The coordinates derived from REST simulations are coloured in green (α4), purple (β2), and grey (ligand). The coordinates from the Cryo-EM structures are coloured in transparent black.*

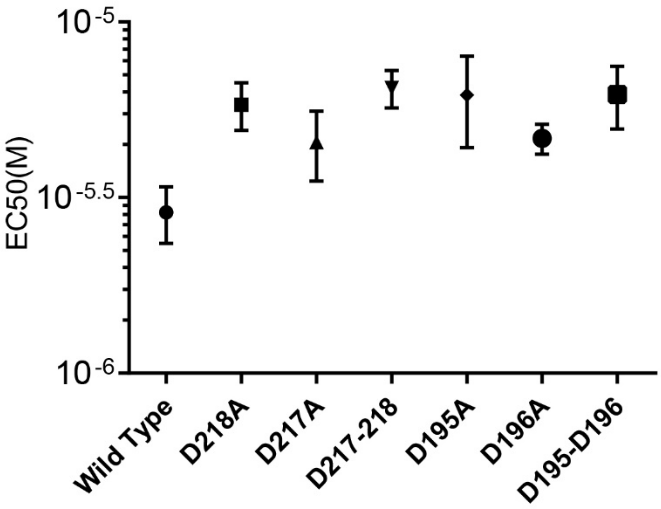


**Figure S2**. *Effect of mutants on the $EC_{50}$ of ACh compared with the wild-type. One-way ANOVA with multiple comparisons showed no significant difference (p-value is 0.1260) between the mutants and the wild-type*.
